# Robust and reproducible neuronal differentiation of human embryonic stem cells for neurotoxicology

**DOI:** 10.1101/2022.01.26.477818

**Authors:** Athina Samara, Martin Falck, Mari Spildrejorde, Magnus Leithaug, Ganesh Acharya, Robert Lyle, Ragnhild Eskeland

**Affiliations:** Division of Clinical Paediatrics, Department of Women’s and Children’s Health, Karolinska Institutet, K6, SE-17177 Stockholm, Sweden; Astrid Lindgren Children’s Hospital Karolinska University Hospital, SE-17177 Stockholm, Sweden; Department of Biosciences, University of Oslo, Blindern, PO Box 1066, 0316, Oslo, Norway; PharmaTox Strategic Research Initiative, Faculty of Mathematics and Natural Sciences, University of Oslo, 0316 Oslo, Norway; Department of Medical Genetics and Norwegian Sequencing Centre, Oslo University Hospital, Kirkeveien 166, Oslo, 0450, Norway; Institute of Clinical Medicine, Faculty of Medicine, University of Oslo, 0450 Oslo, Norway; Division of Obstetrics and Gynecology, Department of Clinical Science, Intervention and Technology (CLINTEC), Karolinska Institutet, Alfred Nobels Allé 8, SE-14152 Stockholm, Sweden; Center for Fetal Medicine, Karolinska University Hospital Huddinge, SE-14186 Stockholm, Sweden; Centre for Fertility and Health, Norwegian Institute of Public Health, PO 222 Skøyen, Oslo, 0213, Norway; Institute of Basic Medical Sciences, Department of Molecular Medicine, University of Oslo, Blindern, PO Box 1112, 0317, Oslo, Norway

## Abstract

Neuronal differentiation from pluripotent cells is commonly used to recapitulate events in early brain development. Neuronal precursors and developing neurons cultured in vivo, rely on attachment, density and cell-to-cell communication, and cell-passaging steps should be minimised to avert disruption of network cross-connectivity. This is crucial both for viability and maturation, and the former also applies to human embryonic stem cells (hESCs). Neuronal differentiation protocols from hESCs for neurotoxicology assessment studies should therefore provide standardized cell density requirements, and optimised coating matrix conditions. The effect of drug treatments may be masked by compound instability and degradation in cell culture medium but performing daily media changes can circumvent these issues. Here, we describe a robust neuronal differentiation protocol using dual SMAD/WNT signalling inhibitors LDN193189, SB431542 and XAV939 (LSX) for neural induction of hESCs, followed by self-patterning and cell maturation stages, for the generation of ventral telencephalic progenitors and neurons. We provide critical information on the optimized cell culture parameters and standardized methods of stage-specific validation. Using cell counts, immunofluorescence, qRT-PCR, and a proof of principle treatment with valproic acid to showcase issues with drug-induced cell toxicity, we facilitate reproducibility of the protocol.

**Graphical abstract:** 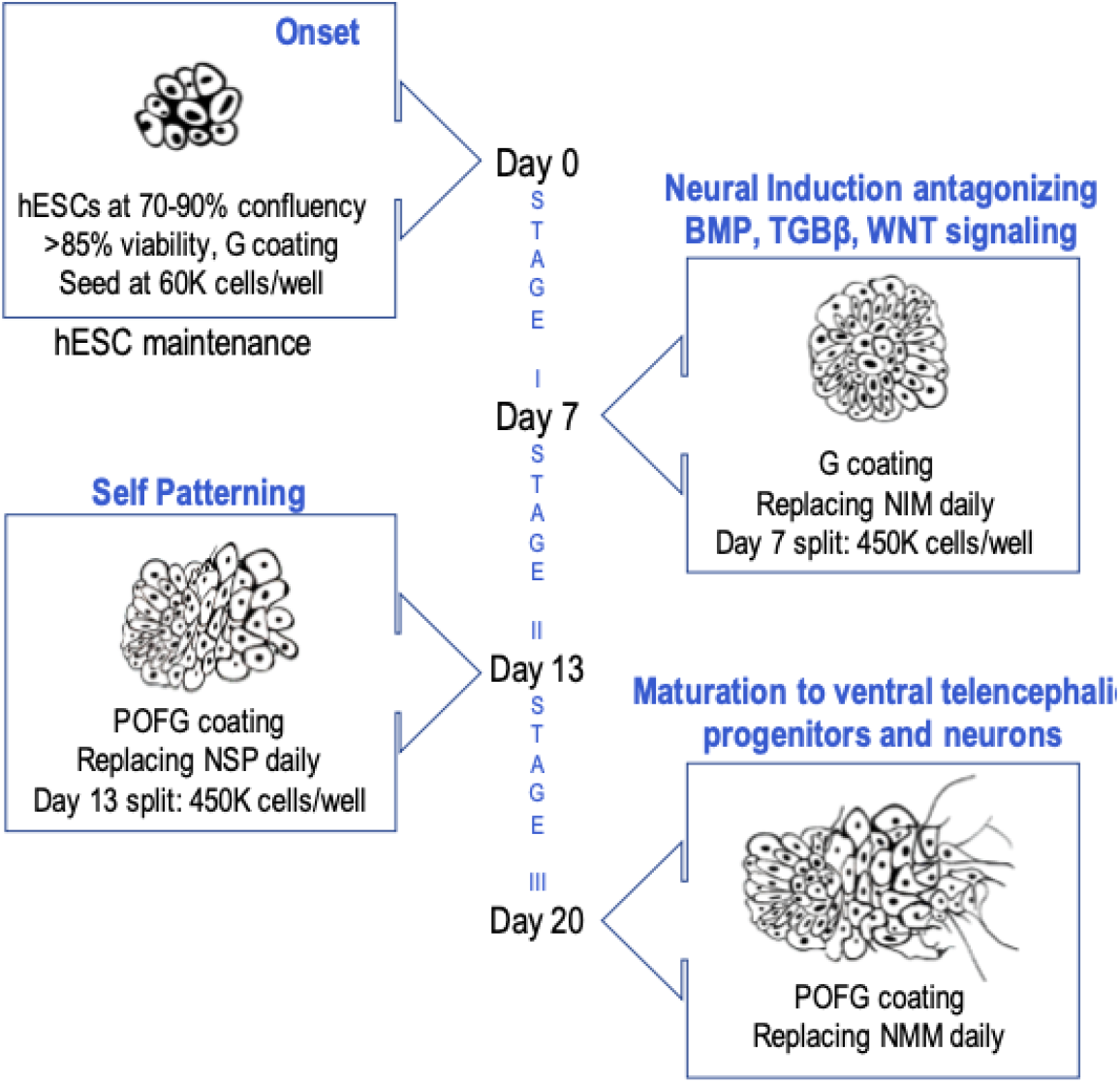

## Introduction

Neural induction of induced pluripotent stem cells or human embryonic cells (hESCs) and differentiation towards targeted neuronal types has wide applicability in neuropharmacology and neurotoxicology studies. Undoubtedly, three-dimensional (3D) culture models offer advantages to the studies of brain architecture, recapitulating the complexity of intraneuronal connectivity. However, for users less experienced with hESCs or neuronal differentiation studies, and studies targeting specific genes or compound effects in early brain development, monolayer approaches offer the advantage of monitoring single cells without the confounding parameters of aggregation seen in 3D cultures. To that end, several protocols have been established for neuronal induction from human pluripotent stem cells and hESCs. A combination of three small molecules (LDN-193189, SB 431542, XAV939) that antagonize the BMP, TGF-ß and WNT signalling pathways, has been documented to drive hESCs to generate cells of forebrain lineage (Ohashi et al. 2018, Cakir et al. 2019, Tchieu et al. 2017, Major et al. 2017).

For toxicology studies of early brain development, the effect of small molecules and drugs may easily be masked, especially at low concentrations. This specific parameter can be addressed with daily media changes to avoid issues of stability and degradation in standard cell culture conditions. Furthermore, after *in vitro* neural induction, serum-free supplements such as N2 and B27 are essential to neuronal differentiation, survival and maturation. The culture conditions should also inhibit the growth of non-neuronal cells (or undifferentiated residual ESCs) and favour the expression of neural specific genes in the derivative cell culture. Well-defined and broadly used coating and culture reagents can also address the effect of drug compounds to proliferation or cytotoxicity, cell adhesion and spatiotemporal effects on the course of differentiation.

Sensitivity to pharmacological treatments vary due to cell type, cell confluency and the availability of space, or cell to cell contact inhibition. Unlike, functional studies of individual neurons that require low density for synapse formation and dendritic spine morphology analyses, neuronal induction protocols should employ a medium-to high-density seeding approach at cell culture splitting, followed by a higher density at replating, to permit optimal cell survival and maturation. Of note, hESCs, neuronal precursors and neurons are contact dependent cells, but contact inhibition is also a known factor affecting cell signalling cascades and gene expression patterns (Ribatti, 2017; Gerard et al, 2014; Carmona-Fontaine et al, 2008). This makes the newly formed cell connections in a monolayer of differentiating cells fragile and susceptible to guidance cues. Thus, cell density at seeding and passaging should be taken under consideration to avoid repetitive mechanical disruption of cell connections due to multiple passaging steps. This is important both for the accurate morphological characterization of the chosen time points of analysis and for the future pharmacological studies (Biffi et al, 2013; Ge et al, 2015).

To this end, we sought to optimise and standardise density and coating parameters and culture conditions to develop a two-split, three stage protocol of neuronal differentiation from hESCs in a monolayer, to study the potential effect of drugs in early forebrain developmental events.

## Materials and Methods

All the culture reagents, chemicals, essential kits and small equipment necessary to reproduce the protocol are listed in Table 1.

**Table 1.**
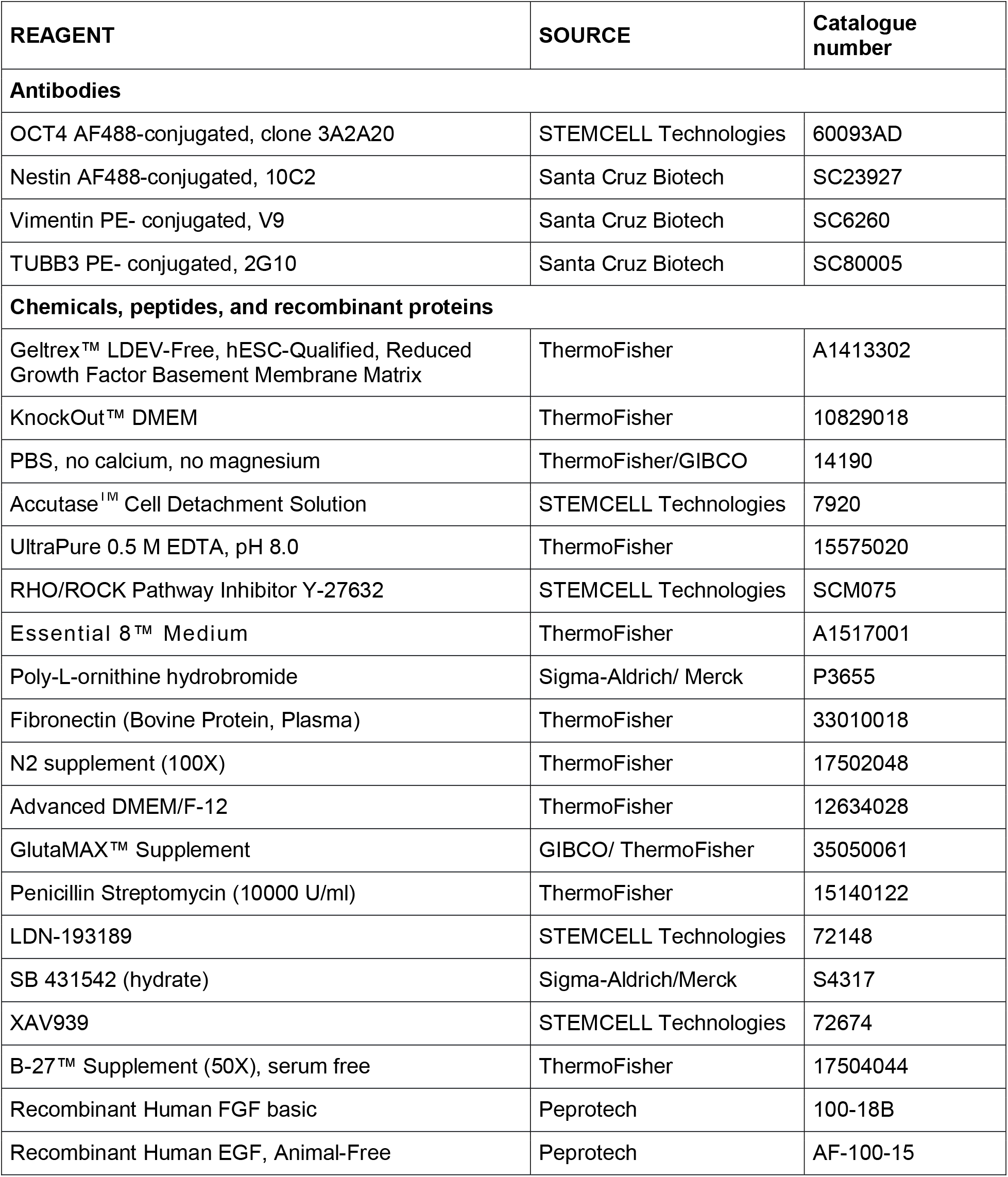
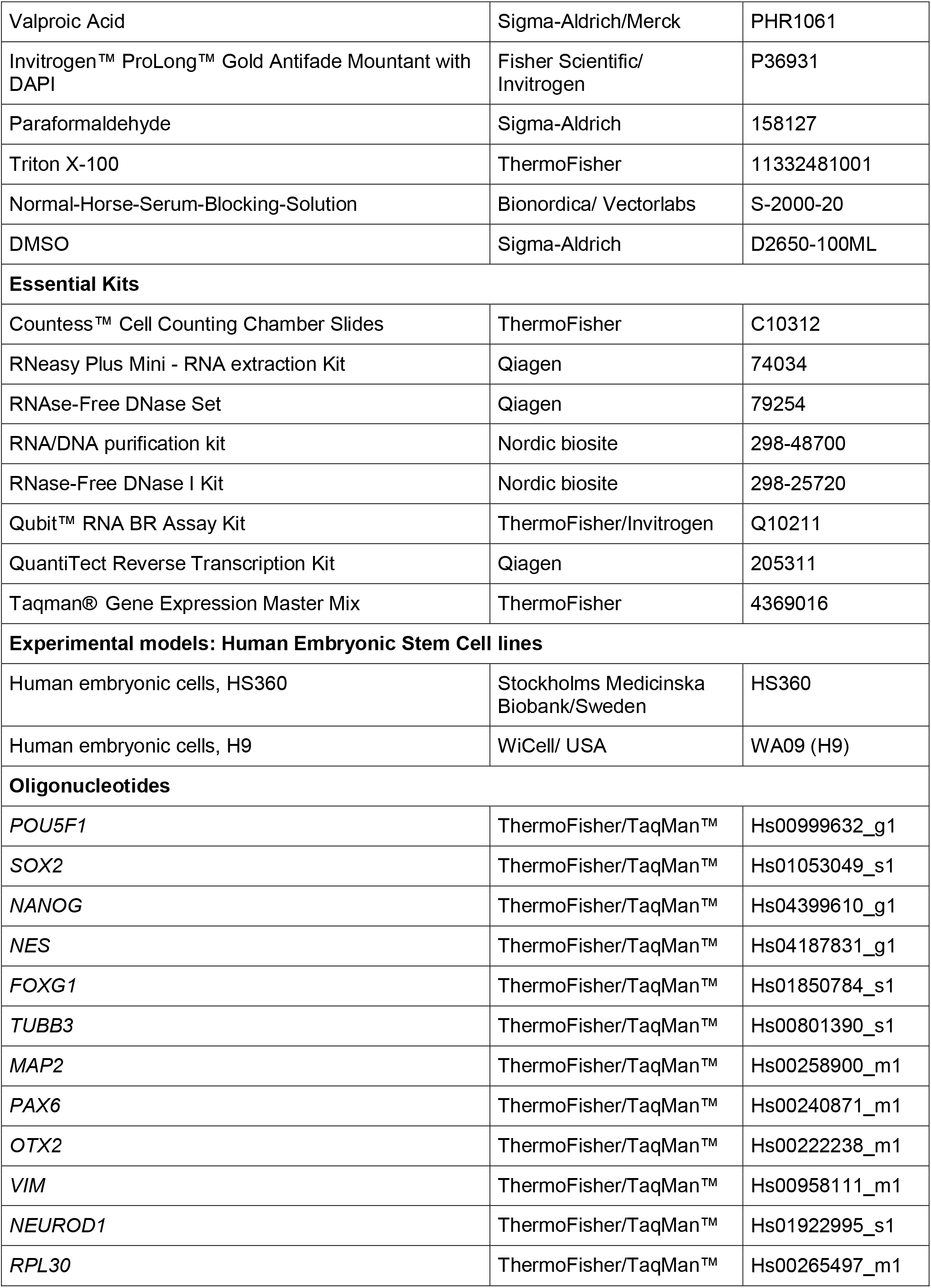
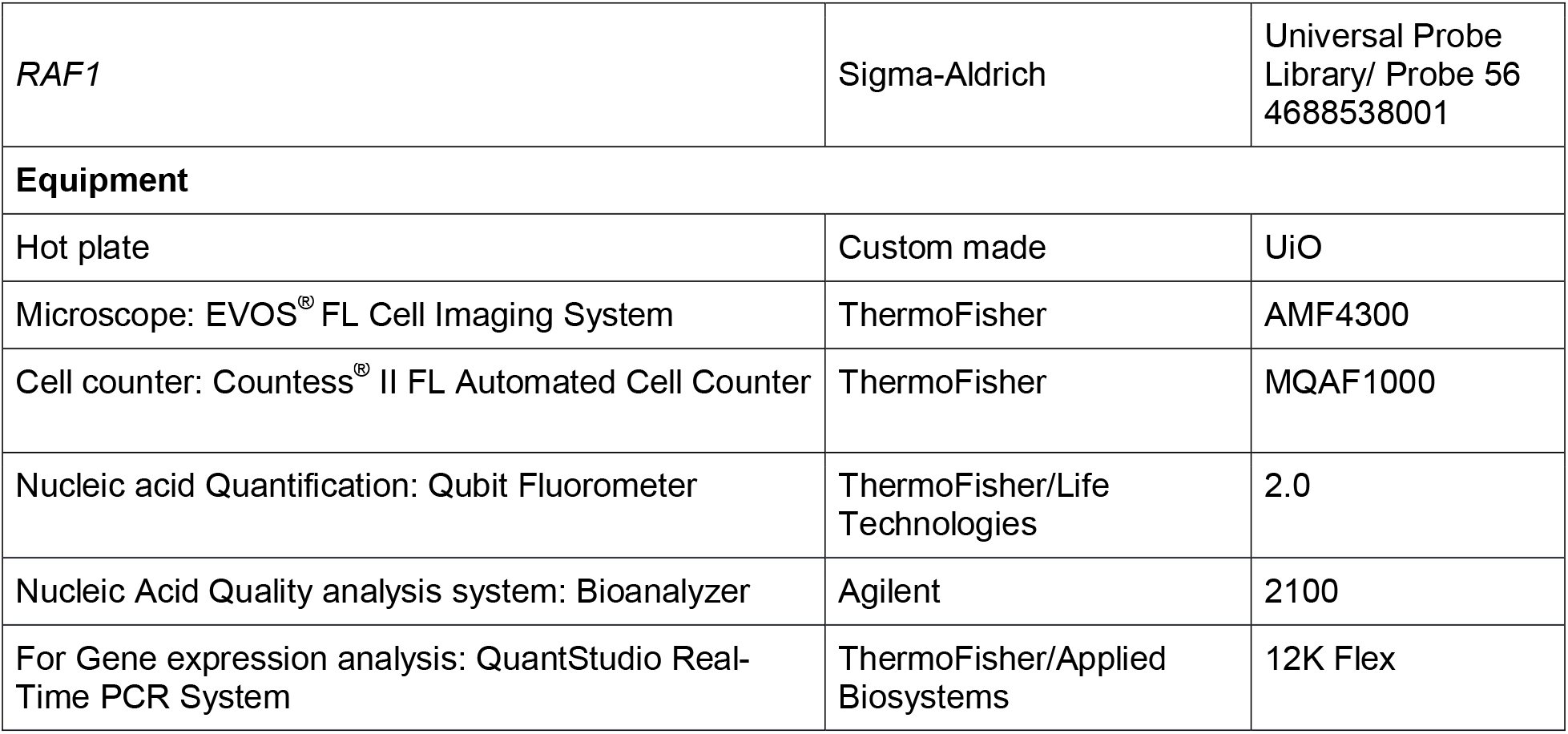
List of reagents and routinely used equipment throughout this protocol.

### Human embryonic stem cell (hESC) culture and maintenance

Human embryonic stem cells HS360 (Stockholms Medicinska Biobank, Sweden, RRID:CVCL C202) (Ström et al, 2010; Main et al, 2020) and H9 (WA09, WiCell) (Thomson et al, 1998) were maintained in Essential 8™ Medium, in feeder-free conditions on Geltrex pre-coated culture 6-well plates (diluted 1:100 in KnockOut DMEM). After thawing hESC stock vials, hESCs were routinely stabilised in culture for 2-3 passages, and a 6-well cell culture plate is regularly used for hESC maintenance. In these conditions, using E8 as the cell culture medium, at 75-90% confluency, over a million HS360 or H9 cells can be seeded from a 6-well plate.

For culture maintenance, cells were routinely passaged at 75-85% confluency using 0.5 mM ethylenediaminetetraacetic acid (EDTA) in ratios between 1:3 to 1:6. For routine culture vessel coating, Geltrex was used at a 1:100 dilution in KnockOut DMEM, according to the manufacturer’s instructions (also described in Table 2). In brief, cells were first washed twice with 1X PBS, treated with 0.5 mM EDTA, for 4 minutes at room temperature, collected with a 5ml serological pipette, spun down at 300 x *g* for 4 minutes and reseeded in fresh culture medium on precoated dishes Geltrex basement matrix. Culture media were replaced daily and if there were signs of differentiation, cells were discarded. Cell cultures were routinely tested for mycoplasma infection. When cells were collected to be stored in liquid nitrogen, they were frozen in culture medium with 10% DMSO.

When hESCs were collected for the differentiation protocol, Accutase was used to detach and dissociate cells instead of EDTA. Cells were counted using the Countess II FL Automated Cell Counter (ThermoFisher). Routine light microscopy was performed using the EVOS FL Cell Imaging System (ThermoFisher) and images were captured with a 1.3 Sony ICX445 monochrome CCD camera (part of the EVOS system).

**Table 2.** Growth factors, cytokine, small molecules, and coating solution stock and working solution preparation.

|  |  |
| --- | --- |
| **SB 431542 (hydrate)** To achieve a final <0.1% DMSO concentration in the culture medium, dissolve SB431542 in DMSO at highest recommended concentration.<br>• Dissolve 5 mg SB431542 in 650 µl DMSO for a stock concentration of 20 mM.<br>• Stock to be aliquoted according to experimental design. For example, if 20 ml of culture media are to be used daily, 10 µl aliquots are prepared and stored at -20°C.<br>• Use a final concentration of 10 µM in culture medium (1:2,000, 0.5 µl/ml media). | **LDN-193189**<br>• Dissolve 10 mg LDN-193189 in 22.6 ml DMSO for a stock concentration of 1 mM.<br>• Place tube(s) in waterbath (37°C), with occasional vortexing until the compound dissolves, making sure there are no undissolved particles.<br>• Stock to be aliquoted in volumes suiting the experimental design (for example, if 20 ml of culture media are to be used daily, 2 µl aliquots are prepared and stored at -20°C.<br>• Use a final concentration of 100 nM in culture medium (1:10,000); dilute once more 1:10 in ddH<sub>2</sub>O for a more convenient 1:1,000 working stock solution. |
| **XAV939**<br>• Dissolve 10 mg XAV in 1.6 ml DMSO for a 20 mM stock concentration.<br>• Aliquot stock and store at -20°C.<br>• Use a final concentration of 2 µM in culture medium (1:10,000); dilute 1:10 in ddH<sub>2</sub>O for a more convenient 1:1,000 work stock. | **RHO/ROCK Pathway inhibitor Y-27632**<br>• Dilute to 10 mM in ddH<sub>2</sub>O<br>• Small aliquots may be stored at -20°C, avoid repeated freeze-thaw cycles.<br>• Use a final concentration of 10µM in culture medium (1:1,000) |
| **Recombinant human FGF basic**<br>Centrifuge vial containing powder prior to opening. Once dissolved, store at 4°C for 1 week, or -20°C to -80°C for long-term storage. Avoid freeze/thaw cycles; recommended storage time in this form and temperature is approximately 12 months.<br>• Reconstitute in 1X PBS 0.1% BSA to 10 µg/ml. Do not vortex.<br>• Aliquot stock solution: e.g., 25 µl in 25 ml of culture medium (if used at 1:1,000).<br>• Use a final concentration of 10 ng/ml in culture medium. | **Recombinant human EGF basic**<br>• Once dissolved, store at 4°C for 1 week, or -20°C to -80°C for long-term storage. Dissolve 1,000 µg in 10 ml of 1X PBS (0.1% BSA added for long-term storage). This results in a 100 µg/ml solution (10,000X stock)<br>• Aliquot 10 ml volume into suitable smaller volumes, for example 100 µl, and store at -20°C.<br>• Use a master/stock aliquot to make a 10X dilution and prepare working solution aliquots. For example, use 100 µl and add 900 µl of 1X PBS (0.1% BSA) to make a 10X 100 µl aliquot 1:1000 working solution.<br>• Use final concentration of 10 ng/ml in culture medium. |
| **Poly-L-ornithine hydrobromide**<br>Prepare in cell culture grade water by dissolving powder to a 10 mg/ml solution, dilute to 0.1 mg/ml with ddH<sub>2</sub>O, and further dilute 1:500 for the stock solution. The stock solution can be stored at 4°C for several weeks, and at -20°C for long-term storage. | **Geltrex basement membrane matrix**<br>• Thaw stock vial in an ice containing beaker containing ice, in a 4°C refrigerator overnight.<br>• Keep tubes on ice while making aliquots.<br>• Ready to use Geltrex working solution is prepared by adding the 250 µl stock in a 50 ml Eppendorf tube containing 24.75 ml of Knockout DMEM. Aliquot 250 µl of working solution in 0.5 ml tubes. |
| **Fibronectin**<br>Dissolve contents of vial in cell culture grade water. Dilute to 1 mg/ml in ddH<sub>2</sub>O by gently warming the vial to 37°C. This is further used as a 1:1,000 stock solution. Do not agitate as it may cause precipitation. Store stock solution at -20°C. | For coating, a stock solution aliquot is thawed on ice or at 4°C and diluted 1:100 in serum-free media. Coat using a volume that will cover the surface of the well/plate (e.g., 1 ml for a well in a 6-well plate); tilt culture plate sideways till surface is covered, making sure there are no dry patches.<br>•Place culture plate in incubator (37°C) for 1 hour, before use.<br>•Remove Geltrex solution and seed cell suspension.<br>Precoating: for long-term storage of coated plates, seal plates with parafilm and store at 4°C for a maximum of 2 weeks. Before use, check that surface is covered and that Geltrex has not solidified or dried out. |

### Preparation of cells and reagents at the start of the protocol

A few important points in the protocol for users less experienced with human embryonic stem cells (hESCs) or neuronal differentiation studies need to be clarified, as reproducibility of the protocol depends on accurate cell counts and high cell viability. Before protocol onset and cell seeding, make sure that there is an abundance of hESC cells in culture that are growing at 80-90% confluency. Stage I Neural Induction Medium (NIM) should be prepared, as described in Table 3. As NIM contains N2 supplement, see manufacturer’s instructions beforehand. If N2 is aliquoted and the penicillin/streptomycin solution thawed, the required time for medium preparation is 10 minutes. Precoat culture dishes with Geltrex for this part of the protocol following coating instructions for cell culture plates for Stage I (Table 4) and see also recipes for preparation of stock solutions as described in Table 2. Timing of preparation is estimated to be 1.5 hours, including coating of culture dishes.

### Critical information for protocol reproducibility

1. Differentiating cells are sensitive to disturbance but rapidly reach confluence. Thus, at daily cell culture medium changes, care should be taken not to disrupt the monolayer. To remove media, the culture dish is lifted from the back at an angle that lets the medium well-up at the front, and the pipettor is placed vertically, to permit medium removal without contact and without disturbing the cells. 80% of the medium is removed, so that the cells do not dry out while replacing media. To replace media, the front edge of the culture dish sits on the hood floor, tilted forward, and medium is changed placing the pipette tip at an angle, by the side at the wall of each well, adding the culture medium dropwise and slowly, protecting the monolayer from mechanical disruption.
2. Cell culture viability might seem low after a single cell count. Thus, as the standard centrifugation conditions suggested are not damaging the differentiating cells, a mild pellet wash in the culture medium the cells will be passaged to, and an additional cell count might be used to clarify the percentage viability of the cells. This is particularly important especially for neurotoxicology experiments.
3. The times for cell detachment using Accutase refer to the use of Accutase prewarmed to 37°C. hESCs in this protocol are maintained in antibiotic free conditions. As hESCs are very sensitive to CO_2_ and temperature changes, they may detach from the culture vessel if these fluctuate. Thus, CO_2_ and temperature should be checked regularly and, if possible, a dedicated cell incubator should be used. Plasticware from different companies has been tested with no effect on cell cultures under the recommended coating conditions. For all centrifugation steps, the temperature was set to room temperature and the conditions (for cell collection and washing) were 300 x *g* for 4 minutes.
4. After cell collection and replating, to ensure even cell spreading, slide the plate back and forth, left and right about 5 times in each direction, avoiding a circular motion which may cause cells to roll back into the centre of wells. Instead, dishes should be moved horizontally (5 times), and side-to-side (5 times), slowly, and carefully moved to the cell incubator.

**Table 3.** Composition of the cell culture media used in Stages I to III of the protocol.

| Composition | Component |  | Amount for 0.5 L total volume |
| --- | --- | --- | --- |
| **Stage I:**<br>**NIM**<br>**(Neural Induction**<br>**Medium)** | **Base**<br>**Medium** | Advanced DMEM/F12 | 485 ml |
|  |  | GlutaMAX Supplement | 5 ml (1%) |
|  |  | Penicillin / Streptomycin | 5 ml (1%) |
|  |  | N2 Supplement | 5 ml (1%) |
|  | SB431542 (added daily) |  | 10 µM final concentration |
|  | LDN-193189 (added daily) |  | 100 nM final concentration |
|  | XAV939 (added daily) |  | 2 µM final concentration |
| **Stage II: NSPM**<br>**(Neuronal Self-**<br>**Patterning**<br>**Medium)** | **Base medium** |  | 495 ml |
|  | B27 Supplement (added to base medium at Day 7 or daily) |  | 5 ml (1%) |
| **Stage III:**<br>**NMM**<br>**(Neuronal**<br>**Maturation**<br>**Medium)** | **Base medium** |  | 497.5 ml |
|  | B27 Supplement (added to base medium at Day 13 or daily) |  | 2.5 ml (0.5%) |
|  | FGF2 (added daily) |  | 10 ng/ml final concentration |
|  | EGF (added daily) |  | 10 ng/ml final concentration |

**Table 4.** Recommended cell culture media, coating, cell detachment and washing volumes, and cell seeding counts. The same conditions apply and have been tested with glass coverslips.

|  | **12 - well format** | **24 - well format** |
| --- | --- | --- |
| **Base matrix coating volume** | 0.5 ml/well | 0.3 ml/well |
| **Cell culture medium volume** | 1 ml/well | 0.5 ml/well |
| **Volume of 1X PBS used for washing** | 1 ml/well | 0.5 ml/well |
| **Volume of Accutase used for cell detachment** | 1 ml/well | 0.5 ml/well |
| **Day 0 cell seed counts** | 60k cells/well | 30k cells/well |
| **Day 7 cell seed counts** | 450K cells/well | 225K cells/well |
| **Day 13 cell seed counts** | 450K cells/well | 225K cells/well |

### Step-by-step method details

#### Stage I: Neural Induction, Time 7 days

##### Day 0 Preparation of culture dishes and media and step by step Stage I instructions

1. At Day 0, ensure that the culture dishes are coated as described, and prewarmed in the incubator. Refer to tables 1–3 for reagents, coating and NIM medium composition instructions.
2. Starting with 80-90% confluency of undifferentiated hESCs, wash cells twice with 1X PBS, and add prewarmed Accutase. The optimal Accutase incubation time for hESC lines HS360 and H9 is 7 minutes.
3. Place culture dish containing Accutase in the incubator or on a hot plate in the hood at 37°C.
4. Protocol efficiency relies on the suspension having a high cell viability at seeding (ideally above 85%). As hESCs are sensitive to vigorous pipetting, the incubation time in Accutase can be extended by 1-2 minutes instead of using mechanical force to attain an easy to collect single-cell suspension.
5. Cells are collected using a 5 ml serological pipette.
6. Single-cell suspension is transferred to Essential 8 medium containing 10 μM ROCK inhibitor (E8/ROCKi), and a small aliquot of the suspension is used for the cell counting.
7. The cells are pelleted at standard centrifugation settings at 300 x *g* for 4 minutes, the supernatant is removed, and the pellet is resuspended again in E8/ROCKi medium.
8. While cells are being centrifuged, cells are counted, and viability assessed.
9. Cells are plated at 60,000 cells per well (see Table 4; this applies when using 12-well culture dishes).
10. Even cell spreading instructions should be followed (see critical information for protocol reproducibility

On Day 1, the E8/ROCKi cell culture medium is replaced with the neural induction medium (NIM) containing the LSX inhibitors. LSX inhibitors are added to the prewarmed (37°C) NIM fresh, before replacing culture medium. NIM medium supplemented with LSX inhibitors is changed daily and times of media changes should be noted and reproduced as accurately as possible throughout the protocol. During pharmacological treatments/toxicology experiments, the compounds of interest are added daily to the medium. Day 7 is the last day of Stage I.

#### Stage II: Neuronal self-patterning, Time 6 days

##### A. Preparation of culture dishes and culture media (3.5 hours)

Refer to Tables 1–4 for reagents, coating and NSPM medium composition instructions, and follow the different coating instructions (POF) for cell culture plates used in Stage II. A solution of polyornithine (20 μg/ml) and fibronectin (1 μg/ml), followed by Geltrex solution (1:100), are sequentially used to coat culture plates both for Stage II and Stage III. In brief, a stock polyornithine aliquot is diluted 1:500 in ddH_2_O, and a stock aliquot of fibronectin is added to that for a final concentration of 1μg/ml fibronectin (for a 50 ml polyornithine solution that would mean adding 50 μl of the fibronectin stock solution). Add the POF mix at the recommended coating volumes (Table 4) and incubate the plates at 37°C for 2 hours. Wash carefully with 1X PBS, add the Geltrex solution and incubate at 37°C for another 1h. The coated culture plates are ready to be used or can be alternatively stored at 4°C for up to two weeks. The composition for the Stage II Neuronal Self Patterning Medium (NSPM) are provided in Table 3. See manufacturer’s instructions for N2 and B27 supplements. If N2 and B27 are aliquoted, and the penicillin/streptomycin solution thawed, the required time for the preparation of the NSPM medium is 10 minutes.

##### B. Step-by-step Stage II instructions

1. At Day 7 ensure that culture dishes for stage II are properly coated and prewarmed in the cell incubator.
2. Carefully remove cell culture medium and place culture dish containing Accutase on a hot plate in the hood, or in the cell incubator
3. Protocol efficiency relies on high viability of the cell suspension at seeding (ideally above 90%). As cells are sensitive to vigorous pipetting, and depending on initial confluency, incubation time in Accutase can be extended by 1-2 minutes instead of using mechanical force to attain an easy to collect single-cell suspension.
4. Cells are collected using a pipettor with 100 ml, 5 ml or 1 ml serological pipettes. At this stage, cells can also be collected with a single channel pipette using a 1 ml tip.
5. Single-cell suspension is transferred to NSP medium containing ROCKi and an aliquot of the suspension is used for the cell count.
6. The cells are pelleted at standard centrifugation settings, the supernatant is removed, and the pellet is resuspended again in NSP/ROCKi medium.
7. While cells are being centrifuged, cell counts, and viability can be assessed.
8. When using the 12-well culture dishes, cells are plated at 450,000 cells per well.
9. Even cell-spreading instructions, as previously described, should be followed.
10. At Days 8-12, NSPM is changed daily (ROCKi is only used when passaging cells at Day 7). During pharmacological treatments/toxicology experiments, the compounds of interest should also be added to the medium daily.

#### Stage III: Neuronal maturation, Time 7 days

##### A. Preparation of culture dishes and culture media (3.5 hours)

The coating instructions for cell culture plates used in Stage III are the same as described previously for the plates used at Stage II.

##### B. Step-by-step Stage III instructions

1. At Day 13 ensure that culture dishes for stage II are properly coated and prewarmed in the cell incubator.
2. Carefully remove cell culture medium and place culture dish containing Accutase on a hot plate in the hood, or in the cell incubator
3. Protocol efficiency relies on high viability of the cell suspension at seeding (ideally above 90%). As cells are sensitive to vigorous pipetting, and depending on initial confluency, incubation time in Accutase can be extended by 1-2 minutes instead of using mechanical force to attain an easy to collect single-cell suspension.
4. Cells are collected using a pipettor with 100 ml, 5 ml or 1 ml serological pipettes. At this stage, cells can also be collected with a single channel pipette using a 1 ml tip.
5. Single-cell suspension is transferred to NMM containing ROCKi and an aliquot of the suspension is used for the cell count.
6. The cells are pelleted at standard centrifugation settings, the supernatant is removed, and the pellet is resuspended again in NM/ROCKi medium.
7. While cells are being centrifuged, cell counts, and viability can be assessed.
8. When using the 12-well culture dishes, cells are plated at 450,000 cells per well.
9. Even cell-spreading instructions, as previously described, should be followed.
10. At days 14-19, NMM is changed daily. ROCKi is only used when passaging cells at Day 13. During pharmacological treatments/toxicology experiments, the compounds of interest should also be added to the medium daily.

### qRT-PCR gene expression analysis

The RNA extraction was performed according to the manufacturer’s instructions (see Table 1 for information regarding the RNA extraction kit and list of probes/oligonucleotides). Reverse transcription (RT) of total RNA was performed using QuantiTect Reverse Transcription Kit according to manufacturer’s instructions (Qiagen). cDNA was then amplified using TaqMan^®^ Gene Expression Master Mix (ThermoFisher Scientific) and Roche or TaqMan gene expression assays for the chosen marker genes. Oligos for *RPL30* and *RAF1* were used as normalization controls. The cycling conditions are described below (Table 5).

**Table 5.** PCR cycling conditions using the primers and probes described in Table 1.

| PCR cycling conditions |  |  |  |
| --- | --- | --- | --- |
| Steps | Temperature | Time | Cycles |
| UNG incubation | 50°C | 2 minutes | 1 |
| Polymerase activation | 95°C | 10 minutes | 1 |
| Denaturation | 95°C | 15 seconds | 40 cycles |
| Annealing/extension | 60°C | 1 minutes |  |
| Hold | 4°C | forever |  |

### Quantification and statistical analysis

The resulting Ct-values were analyzed using the ddCt package v.1.40.0 and visualised using the tidyverse package in R 4.0.4. (Zhang et al. 2021). Statistical comparisons were performed in R using t-test in ggpubr package v.0.4.0 (Alboukadel Kassambara, 2020).

### Immunofluorescence

The immunofluorescence analysis was performed according to the method described in the respective material data sheets of the antibody manufacturers. In brief, cells grown on 13 mm glass coverslips were washed once with 1X PBS and fixed in 4% paraformaldehyde for 15 minutes at room temperature. After washing 3 times with 1X PBS, the cells were permeabilised with 0.3% Triton X-100 in 1X PBS for 15 minutes, washed 3 times with 1X PBS, and blocked with 10% serum for 30 minutes. Then, all conjugated antibodies were diluted 1/500 in 1X PBS containing 0.03% Triton X-100 and coverslips were incubated overnight at 4°C. Next, cells on coverslips were equilibrated at room temperature for 2 hours and washed 3 times (15-minute washes) with 1X PBS, at room temperature. The coverslips were mounted on microscope slides using the ProLong^™^ Gold Antifade Mountant containing DAPI to counterstain the nuclei, and according to the manufacturer’s instructions. All the images shown were obtained using the EVOS FL microscope.

## Results

### Description of the Neural Induction at Stage I

Day 0 is the protocol initiation day, and Day 1 is the neural induction initiation day. The formation and maturation of rosettes and their compaction to neural tube-like structures is the characteristic feature of this part of the protocol. Radial patterning is visible in the cell monolayer from day 3, and by day 6 neural rosettes should be clearly visible (Figure 1). Although some less populated areas may be observed (Figure 1G), mostly at the borders of the culture wells, the cells are highly confluent without viability issues. Rosettes at this stage may be looser (Figure 1H) or reach high compaction (Figure 1I). Day 7 marks the end of the neuronal induction part of the protocol and cells are passaged to the self-patterning Stage II.

**Figure 1.**
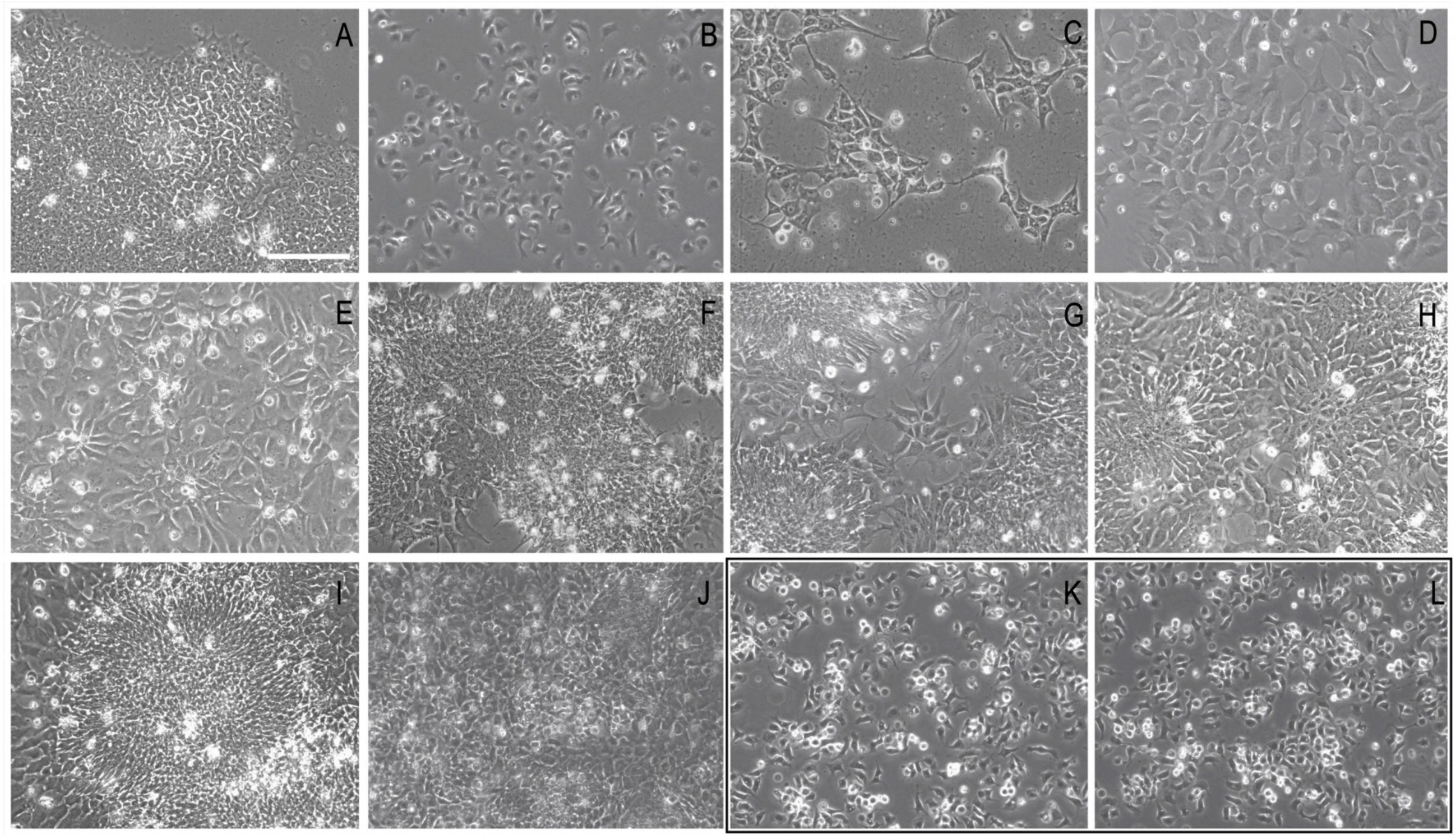
A series of representative brightfield images demonstrating the characteristic formation of rosettes during Stage I neural induction. The protocol starts with the addition of Accutase to the hESC cells in culture (A), and cell counts and seeding for Stage I initiation. (A) Characteristic morphology of the hESCs with the small cytoplasm growing in E8 medium showing no signs of spontaneous differentiation. (B) Day 0, showing the cells 1 hour after plating. (C) Representative image of the culture at Day 1, demonstrating the grid-like formation typical after ROCKi supplementation at Day 1. (D) Proliferation and confluency are enhanced at Day 2 and already at Day 3 the initiation of the columnar arrangement of rosettes can be observed (E). (F-H) The formation and maturation of rosettes and their compaction to neural tube-like structures is the characteristic feature of this part of the protocol. Although some less populated areas may be observed, (as shown at the Day 5 image, (G)), other than few areas at the borders of the culture wells, the cells are highly confluent without viability issues (as assessed by the cell counts). Rosettes at this stage may be looser (H) or reach high compaction (I). (J) At day 7 the cells are collected to be transitioned to Stage II. Images were taken with an EVOS FL microscope at 20X magnification (scale bar corresponds to 100 μm). **Pause point.** Cells can be collected and frozen at this point (Day 7). The black-framed panel shows the cells 1 hour after passaging (K), if the protocol continues uninterrupted or if cells are frozen and reseeded at a later stage (L). Images were taken with an EVOS FL microscope at 20X magnification (scale bar corresponds to 100 μm).

Cells at Day 7 can be collected and frozen (in NIM containing 10% DMSO), to be thawed and reseeded at a later stage. Stage I of the protocol generates cells of anterior neuroectodermal fate using three small molecules (LDN, SB, XAV) that antagonise the BMP, TGF-ß and WNT signalling pathways.

### Stage II is a self-patterning phase

After the Stage I fate induction, this second stage of the differentiation protocol is a growth factor-free, self-patterning stage where the neural precursor monolayer is maintained in N2/B27 supplemented medium. The cells remain adherent whereas survival and maturation is enhanced. Given that the differentiating cells are split and replated twice in 20 days and are plated at high confluency from day 7 onwards, B-27 supplement is added at Stage II to aid neuronal outgrowth and long-term neuronal survival. A set of representative phase contrast images at days 8, 11 and 13 showing how neural stem cells and neuronal precursors organise in the self-patterning stage are shown in Figure 2.

**Figure 2.**
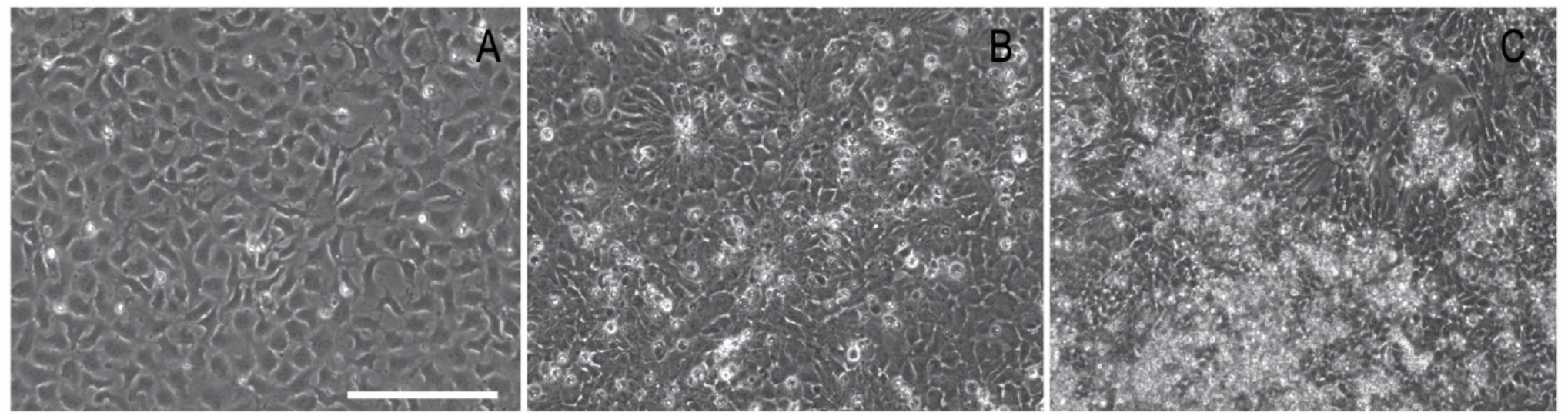
Representative phase contrast images at Day 8, Day 11 and Day 13, showing how the neural stem cells and neuronal precursors (NSCs/NPCs) organise in the self-patterning stage. (A) Image taken at Day 8, the day after plating cells for Stage II. (B) The expanded cells, by Day 11 have the morphology of neural progenitors. (C) By Day 13 the culture forms as a heterogeneous cell population composed of precursors and immature neurons. All images were taken before routine media changes. As described, daily culture medium replacement is not accompanied with washing steps, thus by day 13, dead cells and debris may be accumulating. Images were taken with an EVOS FL microscope at 20X magnification (scale bar corresponds to 100 μm).

### Stage III culture conditions support differentiation and cell maturation

In addition to the N2 supplement and the B27 supplement, which is reduced by 50% compared to Stage II, the Stage III cell culture medium is supplemented with bFGF and EGF to promote growth of maturing neurons. A set of representative phase contrast images at days 14, 17 and 20 showing how the cells organise in maturation stage are shown in Figure 3.

**Figure 3.**
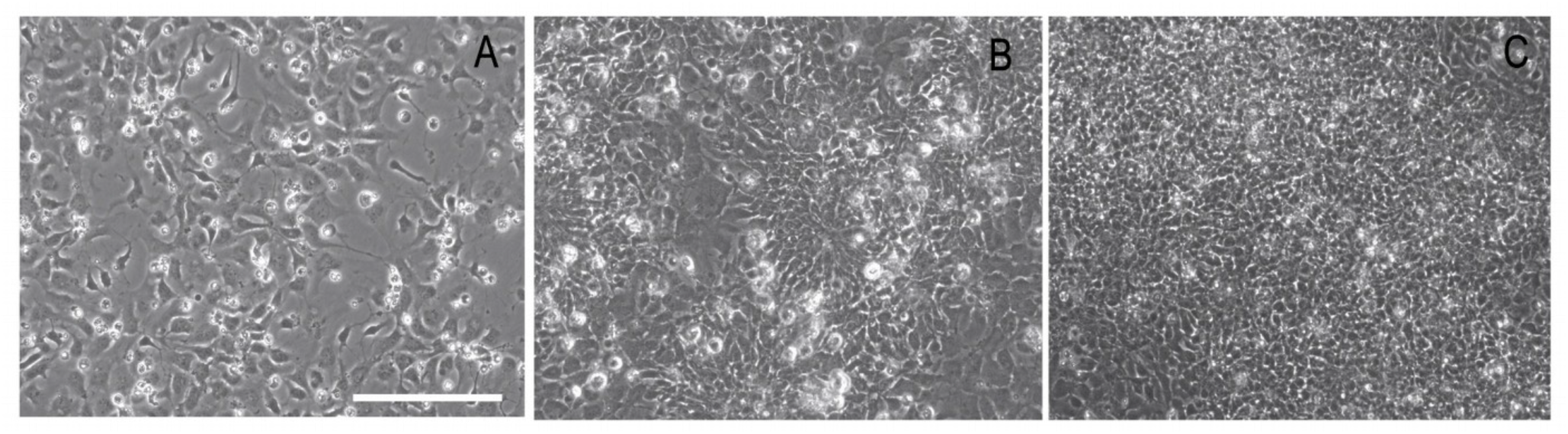
Representative brightfield images at Day 14, Day 17 and Day 20, showing how the culture organises in the maturation stage of the protocol. (A) Image taken at Day 14, the first day after plating cells for Stage III. (B) The cells have, by Day 17 acquired the morphology of expanded NSCs/NPCs. (C). By Day 20 the culture forms a tightly packed and dense small-cell-body population highly reminiscent of a network of NSCs/NPCs and neurons. Images were routinely taken before media changes. As described, daily culture medium replacement is not accompanied with washing steps, thus by Day 20, dead cells and debris may be accumulating. Images were taken with an EVOS FL microscope at 20X magnification (scale bar corresponds to 100 μm).

### Viability and Morphological roadmap of differentiation

Following a protocol is easier when viability reports, cell-counts and visual morphological aids are available. Viability assessment showing cell counts and examples of live/dead reports from the Countess II cell counter at Days 0, 7, 13, 20, from three different protocol replication series are shown in Figure 4.

**Figure 4.**
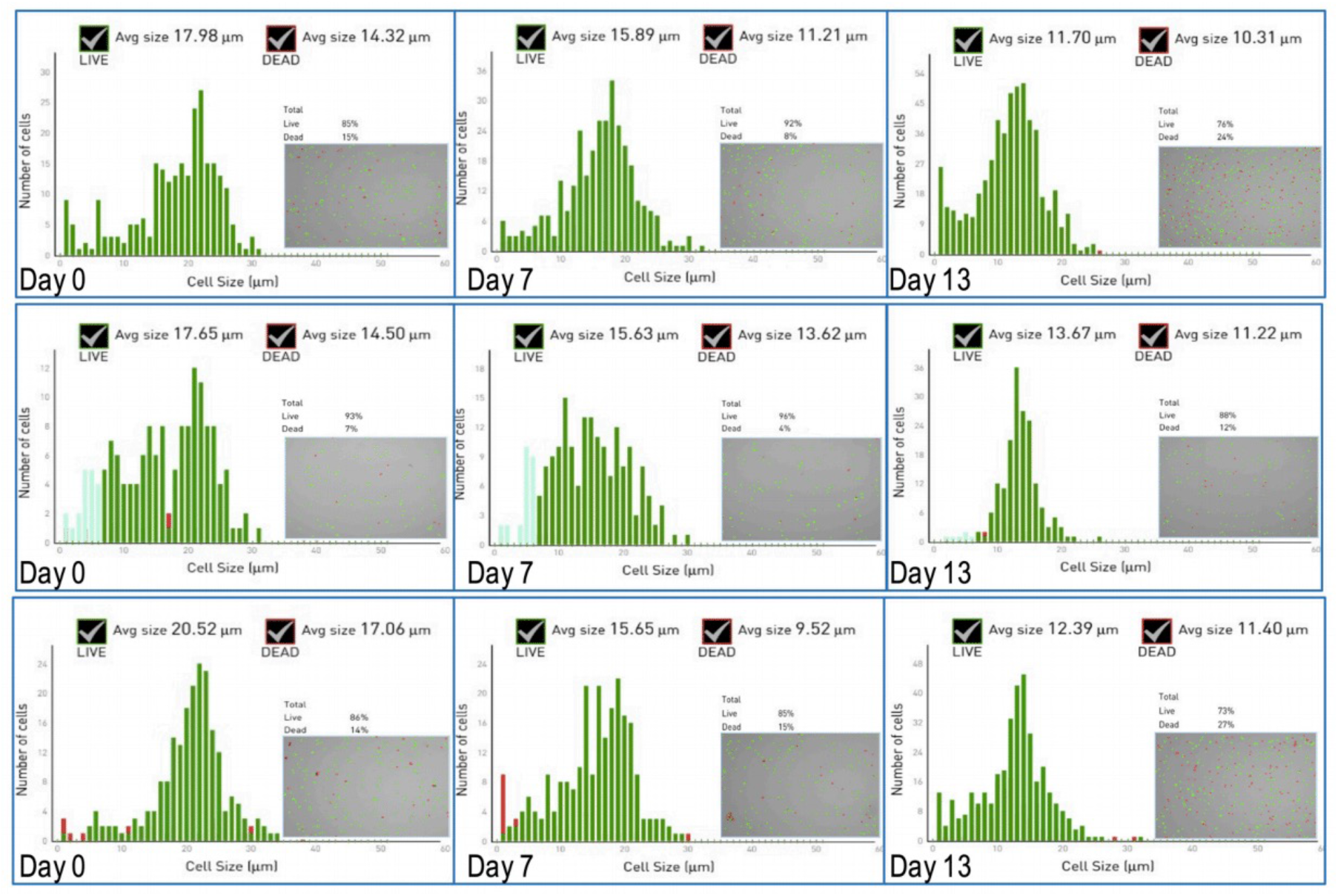
Examples of Countess II Live/Dead Reports of cell counts at Day 0, Day 7 and Day 13 from three different protocol replication series. The raw data from the Countess II readouts are shown as a sequence of 3 graphs. Each report provides information on the cell count, the average cell size of dead (red) and alive cells (green) and the percentage viability as the ratio of live/dead cells. The grey inset in each graph gives more details of the actual cell spread, indicating single cells, doublets and aggregates.

We also provide a series of brightfield images to demonstrate the full timeline of differentiation. As shown in Figure 7, the cells in Stage I start from the confluent hESC cultures at Day 0 and differentiate towards neuroectoderm. Representative images of day-to-day culture reorganization at Stage II (from Day 7 to 13) are shown in Figure 8, whereas images of the cell culture circuit formation from Day 14 to 20 are shown in Figure 9.

**Figure 5.**
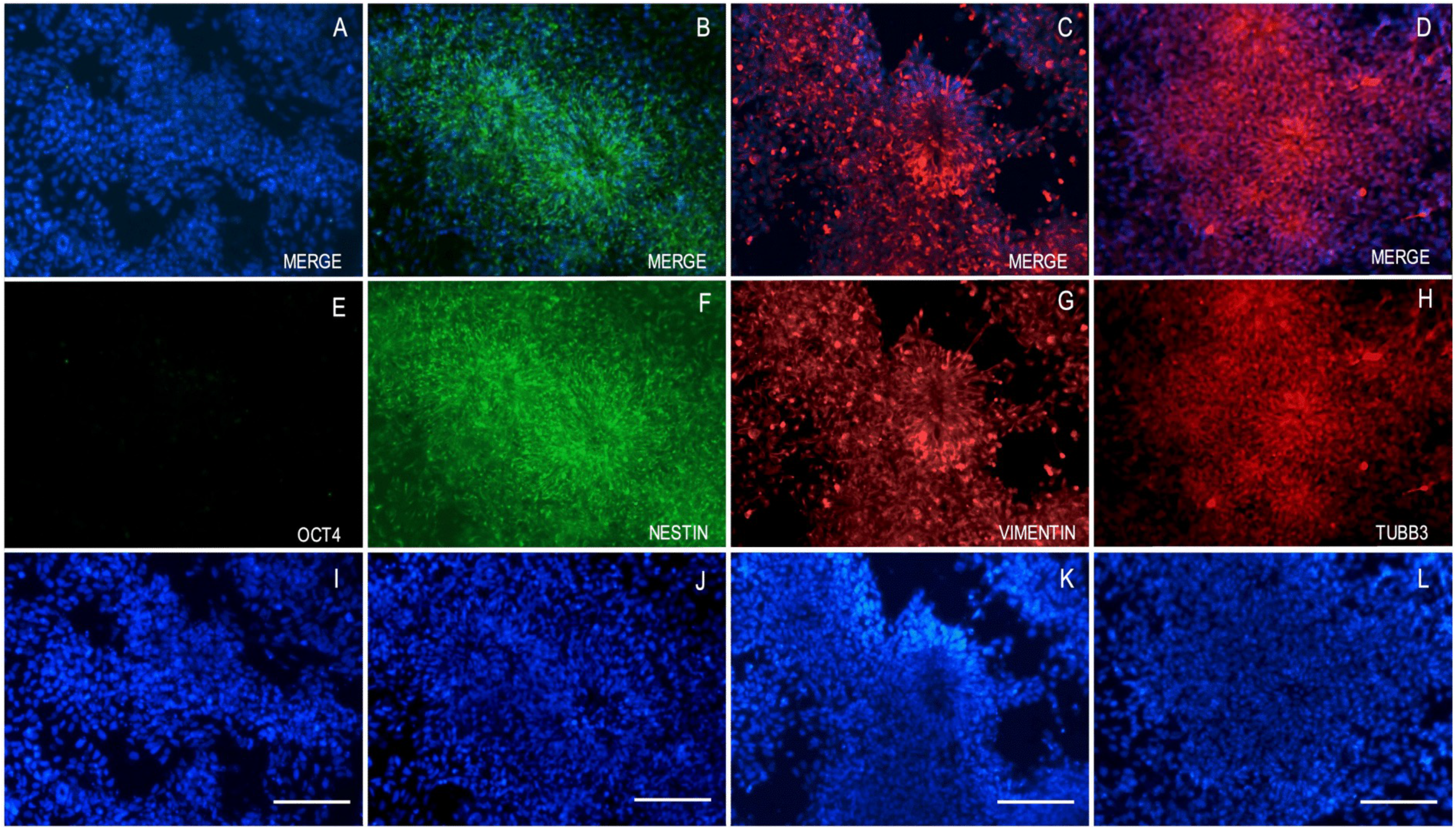
Immunocytochemistry (ICC)/Immunofluorescence analysis at Day 13. To assess the self-patterning stage beyond the characterisation by brightfield microscopy, immunofluorescence imaging analysis was performed at Day 13. A - D are the merged images, with the pluripotency marker OCT4 (E), the intermediate filaments nestin (green, F) and vimentin (red, G) and the neuronal beta-3 tubulin (TUBB3; red, H), respectively, and counterstaining the nuclei with DAPI (blue, I-L). TUBB3 is a cytoskeletal protein expressed in early neurons and thus an immature neuronal marker. The Day 13 differentiated cells were, as expected, devoid of OCT4 expression, while all cells expressed the three cytoskeletal proteins. The images were taken with the EVOS FL microscope, and the scale bar corresponds to 100 μm.

**Figure 6.**
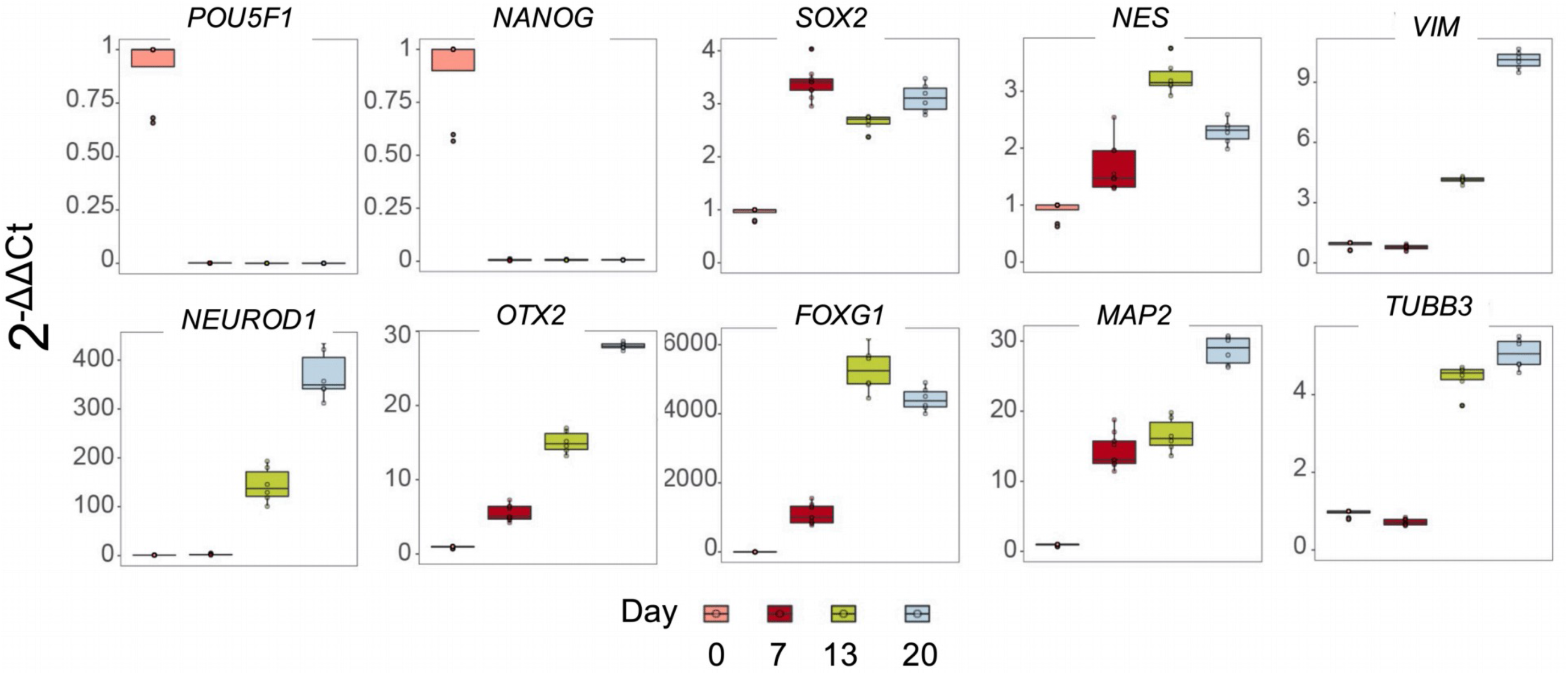
qRT-PCR analysis of pluripotency and neuronal markers. Cells derived at all 3 time points of the differentiation protocol (Day 7, Day 13 and Day 20) were analyzed for specific pluripotency markers (*POU5F1*, *NANOG*, *SOX2*, *NES*), major neuronal development transcription factors (*SOX2*, *OTX2*, *FOXG1*, *NEUROD1*) and genes related to cytoskeletal rearrangement during differentiation towards neuronal maturation (*NES*, *VIM*, *TUBB3* and *MAP2*) by qRT-PCR.

**Figure 7.**
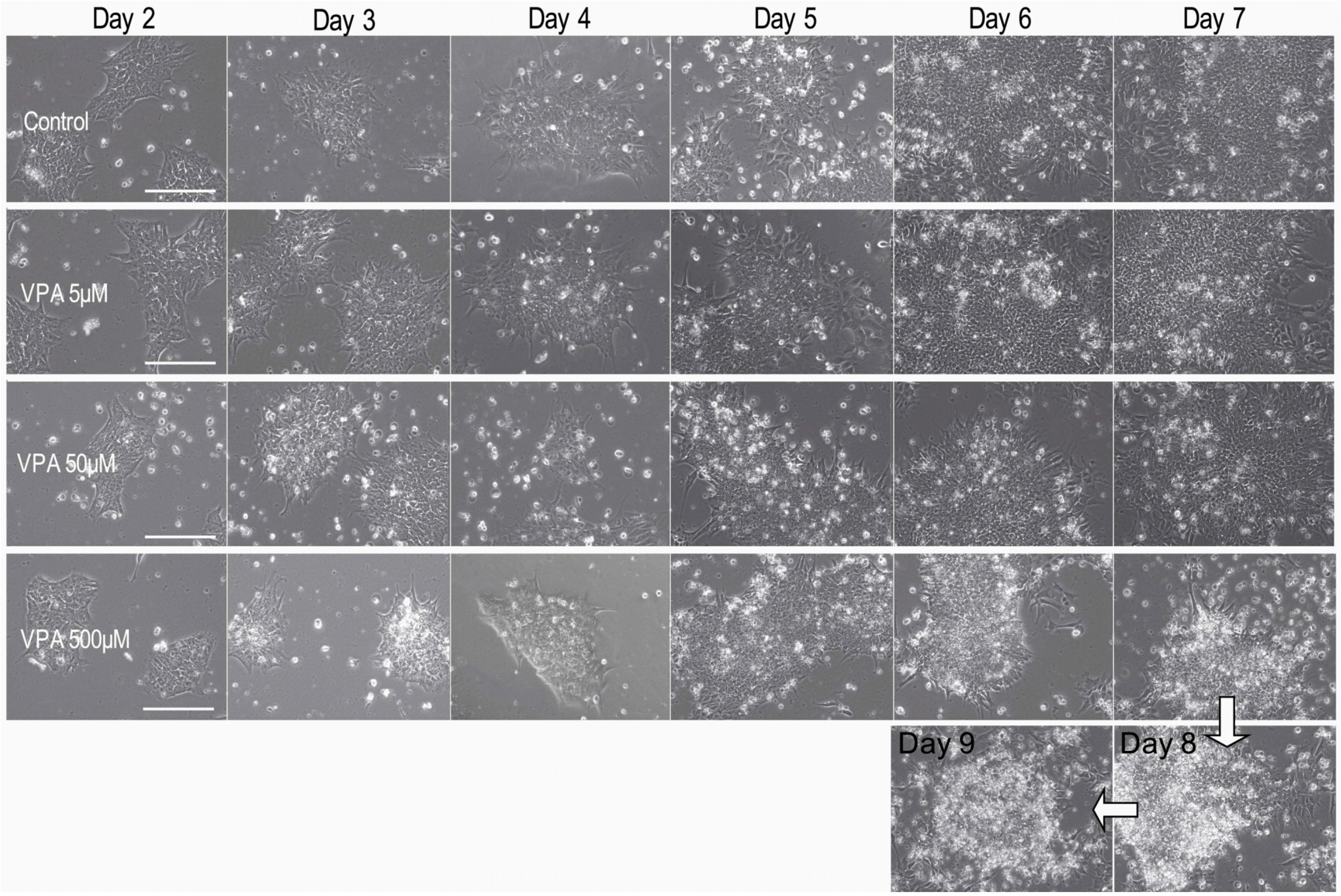
Representative phase contrast images of Stage I of the differentiation timeline in control and VPA treated cells. This detailed roadmap of images of the cells aims to aid the user to reproduce the steps described. As mentioned previously, a high concentration of drug treatment might manifest as increased cell death, with debris and dead cells accumulating on the differentiating live cells. Furthermore, empty or less confluent areas might be more prominent in the culture dish by Day 6. In the example provided, cells treated with 500 μM VPA were maintained in culture for 2 more days, with daily (NIM + 500 μM VPA) media changes, but never reached confluency as seen at the Day 8 and the Day 9 images. This treatment condition was thus paused. The images were taken with the EVOS FL microscope, and the scale bar corresponds to 100 μm.

**Figure 8.**
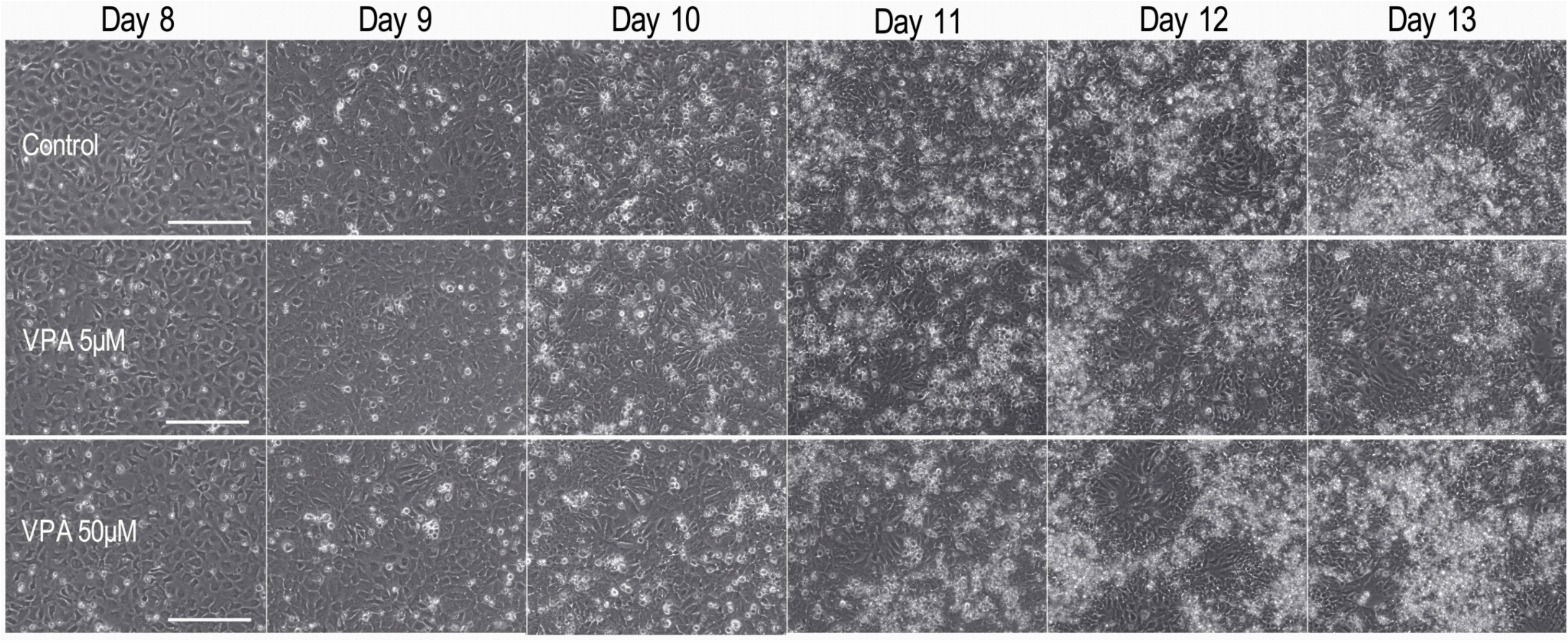
Representative phase contrast images of Stage II (Days 8 to 13) of the differentiation timeline in untreated/control cells. Control, 50 μM and 5 μM VPA treated cells are aligned per day per row. The images were taken with the EVOS FL microscope, and the scale bar corresponds to 100 μm.

**Figure 9.**
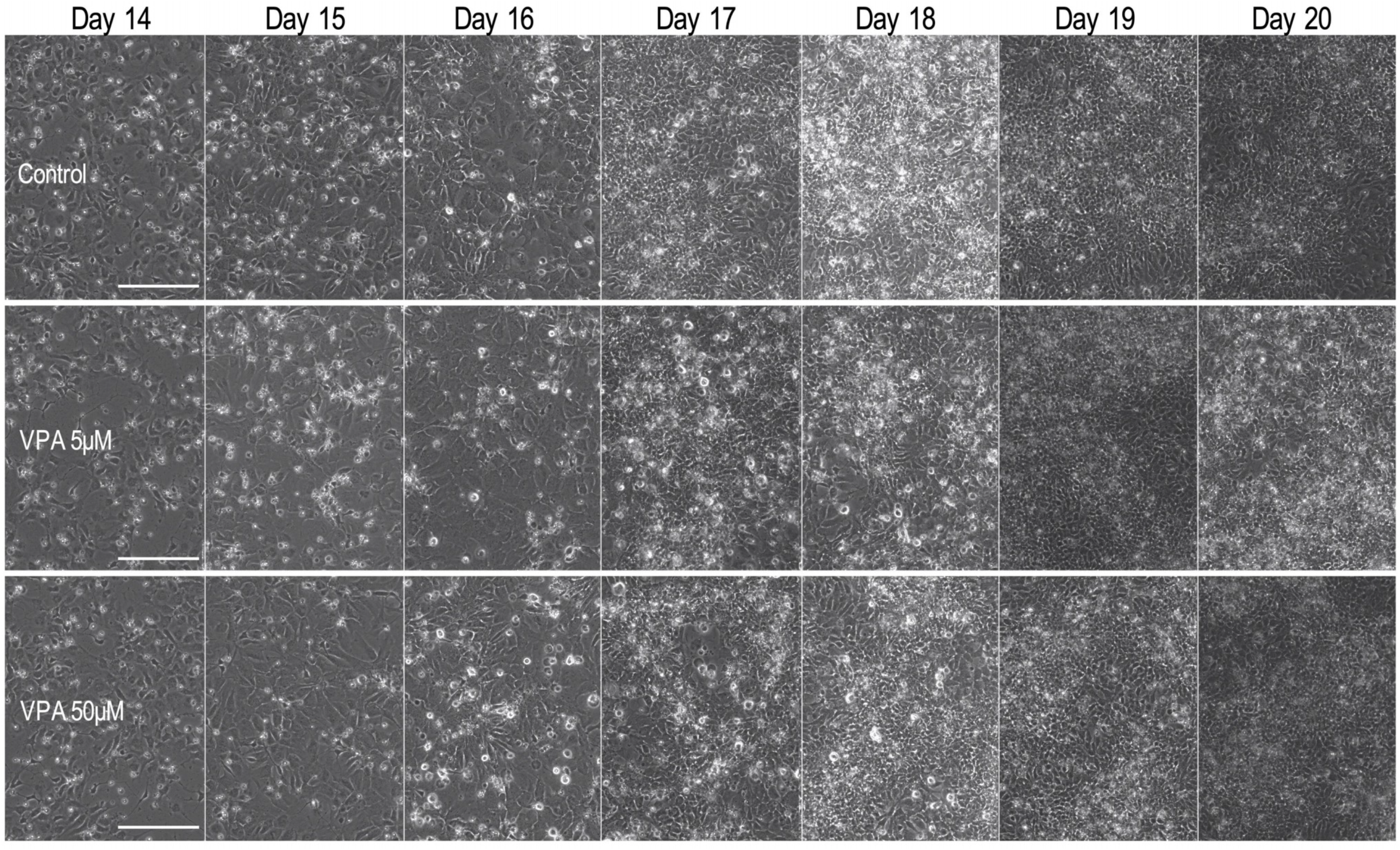
Representative phase contrast images of Stage III (Days 14 to 20) of the differentiation timeline. Control, 50 μM and 5 μM VPA treated cells are aligned per day per row. The images were taken with the EVOS FL microscope, and the scale bar corresponds to 100 μm.

As this protocol was replicated with H9 hESCs we also provide the full morphological timeline of differentiation for these cells in Supplementary Figure 1.

### qRT-PCR analysis of pluripotency and neuronal markers

Gene expression analysis of marker genes as a proxy to follow neuronal differentiation at selected timepoints is a useful and inexpensive tool available locally to most labs. For user convenience we also provide indicative concentrations of RNA and DNA after nucleic acid isolation at Days 0, 7, 13 and 20 (Table 6).

**Table 6.** Indicative RNA and DNA concentrations after the respective nucleic acid isolation and extraction at days 0, 7, 13 and 20.

| Day | DNA yield (µg) | RNA yield (µg) |
| --- | --- | --- |
| 0 (6-well plate) | 1-4 | 21-38 |
| 7 (12-well plate) | 2-5 | 10-17 |
| 13 (12-well plate) | 2-5 | 7-17 |
| 20 (12-well plate) | 2-8 | 11-24 |

In order to characterise the derivative cell populations, hESCs collected at the onset of the protocol (Day 0), and cells derived at all 3 time points of the differentiation protocol (i.e., at Day 7, Day 13 and Day 20) were analyzed for specific pluripotency markers (*POU5F1*, *NANOG*, *SOX2*, *NES*), major neuronal development transcription factors (*SOX2*, *OTX2*, *FOXG1*, *NEUROD1*) and genes related to cytoskeletal rearrangement during differentiation towards neuronal maturation (*NES*, *VIM*, *TUBB3* and *MAP2*) by qRT-PCR.

The expression of the pluripotency transcription factors *POU5F1* and *NANOG* plummets to zero already at Day 7, while *SOX2* and *NES* expression increases, as expected, as hESCs commit to neuronal fate. *NES* expression drops at Day 20, representative of the arising neuronal, non-proliferative cell population of Day 20. The same expression pattern was observed with *FOXG1*, one of the earliest telencephalic specific transcription factors. The expression of the transcription factor *OTX2*, which is known as a regulator of neurogenesis, antagonises ground state pluripotency and also increases as cells differentiate, promoting their commitment. This increasing pattern of expression from Day 0 to Day 20 is also seen in the expression of the key bHLH enhancer of transcriptional regulators of neurogenesis *NEUROD1*, the mature dendritic marker *MAP2* and the neuronal specific *TUBB3*, and the intermediate filament *VIM*.

### Using VPA as a proof of principle pharmacological treatment with a lag phase

A proof of principle example of a pharmacological treatment, using three different concentrations of valproic acid (VPA; 5 μM, 50 μM and 500 μM) and phase contrast images in comparison to control cells are shown (Figure 7, 8 and 9). The Countess II stage-to-stage passaging reports (Figure 10) and the corresponding qRT-PCR analysis of gene expression of the same pluripotency markers, major neural and neuronal development transcription factors and genes related to cytoskeletal rearrangement during differentiation towards neuronal maturation, are also made available (Figure 11). A possible solution, before deciding against using a high drug concentration, would be to extend Stage I for a few days monitoring whether the cells under treatment catch up compared to the indicated neural induction endpoint, as seen at the control cell cultures (Figure 7 and Supplementary Figure 2).

**Figure 10.**
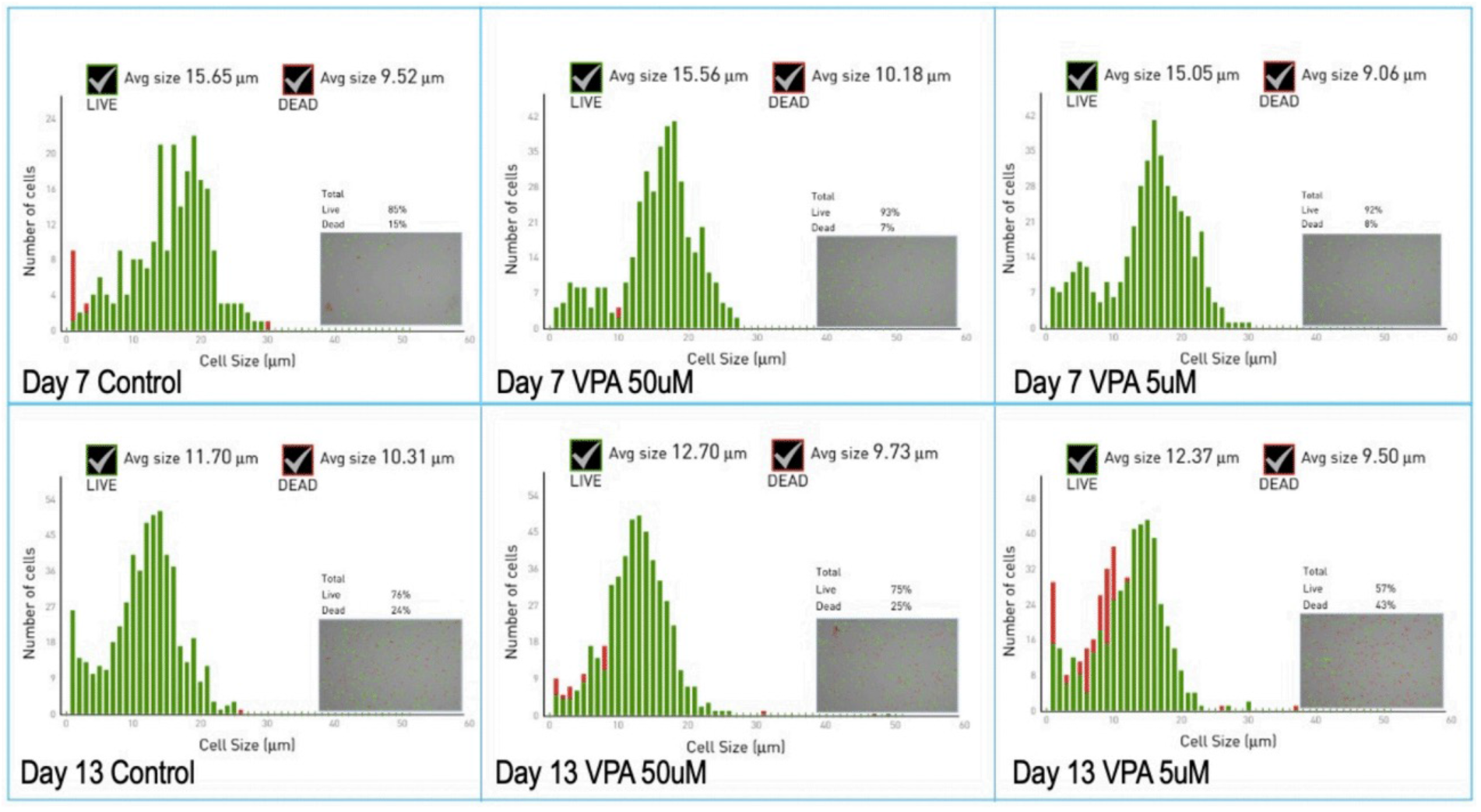
Raw data from the Countess II readouts for the cell counts at Day 7 and Day 13 for the untreated/control cells and the cells treated with 5 or 50 μM of VPA. The counts are shown as a sequence of 3 graphs, showing the Day 7 series on the top, and the Day 13 series of counts below. Each report, as generated by the Countess II provides information on the cell number, average cell size of dead (red) and alive cells (green) and the percentage viability as the ratio of live/dead cells. The grey inset in each graph provides extra information of the actual cell spread, indicating single cells, doublets and aggregates.

**Figure 11.**
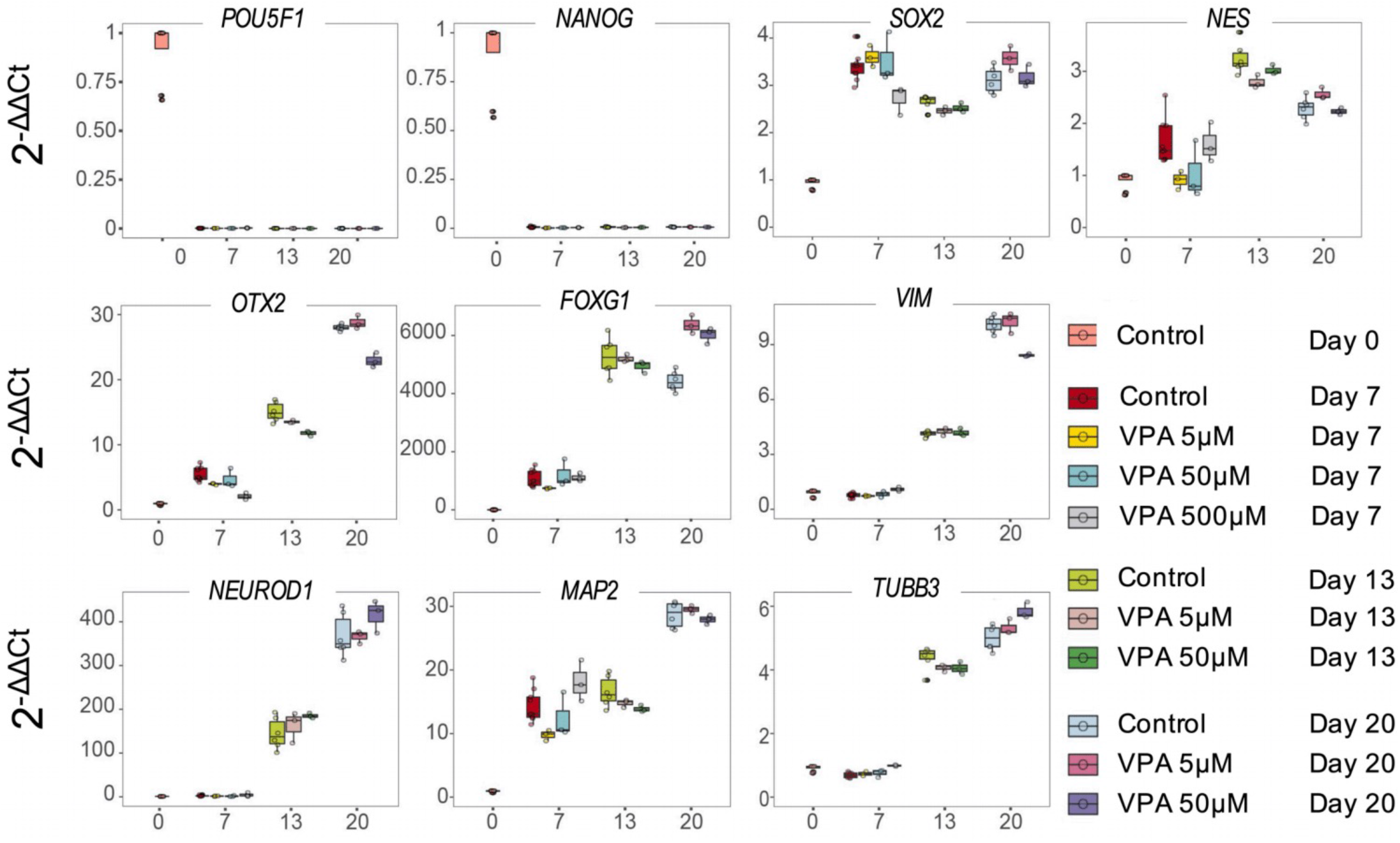
qRT-PCR analysis after VPA treatment. The same marker genes chosen to characterise the derivative cells of the new differentiation protocol, were used to compare the control and VPA treated cells’ expression levels, at all 3 time points (Day 7, Day 13 and Day 20). A set of 3 samples per experiment, from 3 separate experiments were used per treatment, and each sample was run in technical triplicate. Expression levels were analyzed for *POU5F1*, *NANOG*, *SOX2*, *NES*, *SOX2*, *OTX2*, *FOXG1*, *NEUROD1*, *VIM*, *TUBB3* and *MAP2* by qRT-PCR (see methods section for normalization of gene expression levels). The expression levels of all genes at the hESC level (Day 0) are also illustrated for comparison.

### qRT-PCR analysis after VPA treatment

The same marker genes chosen to characterise the derivative cells of the new differentiation protocol presented in Figure 6, were used to compare the control and VPA treated cells’ expression levels, at all 3 time points (Day 7, Day 13 and Day 20). A set of 3 samples per experiment, from 3 separate experiments were used per treatment, and each sample was run in technical triplicate. Expression levels were analyzed for *POU5F1*, *NANOG*, *SOX2*, *NES*, *SOX2*, *OTX2*, *FOXG1*, *NEUROD1*, *VIM*, *TUBB3* and *MAP2* by qRT-PCR. The expression levels of all genes at the hESC level (Day 0) are also illustrated for comparison.

As described previously (Figure 7), the VPA 500 μM treatment was interrupted after Day 7, thus only the results for the Stage I neuronal induction supplemented with 500 μM VPA are available and shown in the qRT-PCR differentiation timeline. The expression of the pluripotency transcription factors *POU5F1* and *NANOG* dropped to zero in all culture conditions and all treatments by Day 7, with subsequent and significant increase of *NEUROD1*, *SOX2* and *NES* indicative of efficient neural induction.

When the culture medium was supplemented with 500 μM from Day 1 to Day 7 (i.e., throughout the neural induction stage), expression of *OTX2* (p < 0.0001) and *SOX2* (p < 0.05) were lower compared to the control. Expression of *MAP2* was slightly increased, though not significantly, whereas the filamentous markers *VIM* (p < 0.05) and *TUBB3* (p < 0.0001) were significantly increased. These differences may be attributed to the increased appearance of larger cells on the periphery of the colonies, in the absence of contact inhibition. The expression of the filamentous marker *NES* did not change significantly in cells exposed to 500 μM VPA compared to control cells at Day 7. In this preliminary analysis, supplementation of the NIM with 500 μM VPA neither enhanced neural differentiation nor exerted a neuroprotective effect.

Supplementation and daily changes of the culture media with 5 μM VPA, apart from a significant increase in the expression of *FOXG1* (p < 0.001) and *NES* (p < 0.05) at Day 20, exerted no significant changes in the expression of the marker genes. Furthermore, exposure to 50 μM VPA, significantly reduced the expression of *OTX2* (p < 0.05) and *VIM* (p < 0.001), and similarly to the 5 μM VPA, exposure to 50 μM VPA significantly increased the expression levels of *FOXG1* (p < 0.001). Additionally, the expression of *TUBB3* (p < 0.01) and *PAX6* (p < 0.001) were significantly increased in cells exposed to 50 μM VPA compared to control cells. Finally, the expression of *NEUROD1* in cells exposed to 50 μM VPA at Day 20, was slightly but not significantly increased when compared to the control cells, or the cells exposed to 5 μM VPA for 20 days.

## Discussion

Here we describe a novel neuronal differentiation protocol using dual SMAD/WNT signalling inhibitors LDN193189, SB431542 and XAV939 (LSX) for neural induction of human embryonic stem cells in a monolayer, followed by a self-patterning and a cell maturation stage.

The novel neuronal differentiation protocol describes the specific steps for using the hESC cell line HS360, and it has been replicated using H9 hESCs. It consists of three major stages: neural induction, self-patterning and neuronal maturation. Cell counts are standardised at Day 0 seeding and the culture medium is changed daily. The cell cultures are split and reseeded at standardised cell numbers in differently coated culture dishes at Day 7 and Day 13, and the protocol ends at Day 20. The same format of culture dishes is used throughout.

We provide a step-by-step description of the protocol, cell count and viability reports, visual aids, such as phase contrast images of the differentiating cells. The protocol is reproducible and robust, and a less costly alternative compared to other similar protocols. It has been optimised for use with widely available coating agents and matrices, previously tested neuronal induction reagents, and standardised cell numbers. In addition, cell numbers and substrate-coating conditions have been optimised to minimise cell passaging and thus to avert mechanical disruption of the cell connections in the monolayer after cell collection. We provide detailed recipes for the preparation of the stock and working solutions, and for the composition of the media used at all stages. In addition, the recommended volumes for the coating of the culture plates, cell detachment and washing, along with cell culture media volumes and cell seeding counts are also made available. The daily media changes suggested, circumvent the possible bias conferred on neurotoxicology experiments by compound instability and degradation in cell culture conditions, which could mask the effect of exposure to the compounds of interest.

We are also sharing information on viability and average cell size both for dead and live cells, as provided by the Countess II reports, and critical information for passaging cells. Furthermore, for reproducibility purposes we have documented the end of the self-patterning stage (Stage II), with representative immunofluorescence images of the cell cultures. Furthermore, we are providing indicative information on the RNA and DNA yield per time point, and the qRT-PCR analysis results of pluripotency markers and major neuronal development transcription factors and genes related to cytoskeletal rearrangement towards neuronal maturation for the timepoints Day 0, 7, 13 and 20. The marker genes chosen were *POU5F1*, *NANOG*, *SOX2*, *NES*, *VIM*, *NEUROD1*, *OTX2*, *FOXG1*, *MAP2* and *TUBB3*.

Based on morphological criteria, the cell network formation Day 20 displayed neuronal characteristics. However, the derivative NPCs and/or neurons generated by this protocol were not assessed for their membrane electrochemical maturation properties, or secretion of neurotransmitters, as it was beyond the scope of this study. This protocol was designed for neurotoxicity assessments aiming at facilitating studies of early brain development events.

The experimental design uses a 12-well culture plate format, allowing for smaller volumes of media, but still providing enough cells for RNA or protein isolation. In a 12-well format, 4 biological triplicates can be tested per plate (for example a triplicate of wells treated under control conditions and 3 triplicates of 3 different doses of the drug treatment of choice). 24-well plate format volumes and cell seeding counts are also described to downscale costs or to use in pilot experiments. Additionally, 13 mm glass coverslips can be placed in the 24-well dishes facilitating immunofluorescence or other type of or microscopy experiments. However, upscaling to 6-well plates might require some optimization beyond the obvious surface area conversion.

hESCs are sensitive to high drug concentrations and neuronal induction is a phenomenon that may induce apoptosis. High drug concentration may thus result in excess cell death manifesting with floating cell debris due to cell death at various stages of the protocol. It may also cause a major lag in the neural induction stage of the protocol (Stage I), and the cells might not follow the self-patterning stage (Stage II). We tested the applicability of the novel protocol with a proof of principle example of such a pharmacological treatment, using increasing concentrations of valproic acid (VPA; 5 μM, 50 μM and 500 μM). VPA was evaluated in this protocol for several reasons. The effect of VPA has been assessed in various studies of multipotent and pluripotent cells towards neuronal differentiation, however the biological mechanism and pathways have not been identified (Duan et al, 2019; Fernandes et al 2019). It has also been documented that VPA, as a histone deacetylase inhibitor, may create a neuroprotective niche (Dekker et al, 2021). Moreover, the effect of VPA *in utero* has also been studied (Ornoy, 2009; Mansour et al, 2021; Zhao et al, 2019) and it was shown that VPA may induce neural tube defects when taken during pregnancy. Of note, neural tube defects have been correlated to increases in the risk of incidence of autism spectrum disorder in children (Arndt et al, 2005; Christensen et al, 2013; Hasler et al 2021). The levels of VPA in the serum of treated patients were 0.2–0.6 mM, but concerns were raised that foetuses might be exposed to higher levels, and up to five times higher than maternal serum at term (Ornoy, 2009). Thus, this protocol was tested as a platform to recapitulate the events at and following neural induction, in neuronal morphology and to delineate these concerns. We showed that before deciding against using a high drug concentration, the protocol may be modified by extension of Stage I for a few days monitoring whether the cells under treatment catch up compared to the indicated neural induction endpoint, as seen at the control cell cultures. The qRT-PCR analysis showed that 50mM VPA induces a statistically significant increase of forebrain markers *FOXG1* and *OTX2* gene expression, but analysis of the molecular cues is beyond the scope of the preliminary VPA analysis shown here.

In conclusion, we present a step-by-step description of a robust and standard characterised protocol for the generation of ventral telencephalic progenitors and neurons. We have highlighted the points in the protocol that are crucial to maintain high cell viability for users less experienced with hESCs or neuronal differentiation studies, to facilitate reproducibility of the protocol. Providing visual aids and gene expression analyses examples for specific marker genes for all timepoints, will aid users that target specific genes or compound effects in early brain development neuropharmacology studies.

## Acknowledgments

Human ES cell line H9 (James A. Thomson, University of Wisconsin) was provided by WiCell Research Institute Inc. We thank Kristina Gervin and Ankush Sharma for fruitful discussions and comments on the manuscript. We acknowledge funding from the Swedish Research Council 2019-01157 (A.S.) and grants from the Swedish Brain FO2019-0087 (A.S.) and the Freemasons Children’s House of Stockholm (A.S.), the Research Council of Norway, 241117 (R.L.); Anders Jahre Foundation (R.E.), Nansen Foundation (R.E.), Wedel Jarlsberg Foundation (M.F.). PharmaTox Strategic Research Initiative was supported by Mathematical and Natural Science Faculty, University of Oslo.

## Author contributions

Conceptualization, A.S., M.F.; Methodology, A.S., M.F., M.S., M.L.; Writing – Original Draft, A.S. M.F.; Writing – Review & Editing, A.S., M.S., R.L., R.E., M.L., M.F., G.A.; Investigation and Validation, A.S., M.F., M.S., M.L.; Funding, A.S., M.F. and R.E.; Supervision, R.L., A.S., and R.E.; Resources, A.S., G.A., R.L. and R.E.

