## Supplementary Figures for "Robust and reproducible neuronal differentiation of human embryonic stem cells for neurotoxicology"

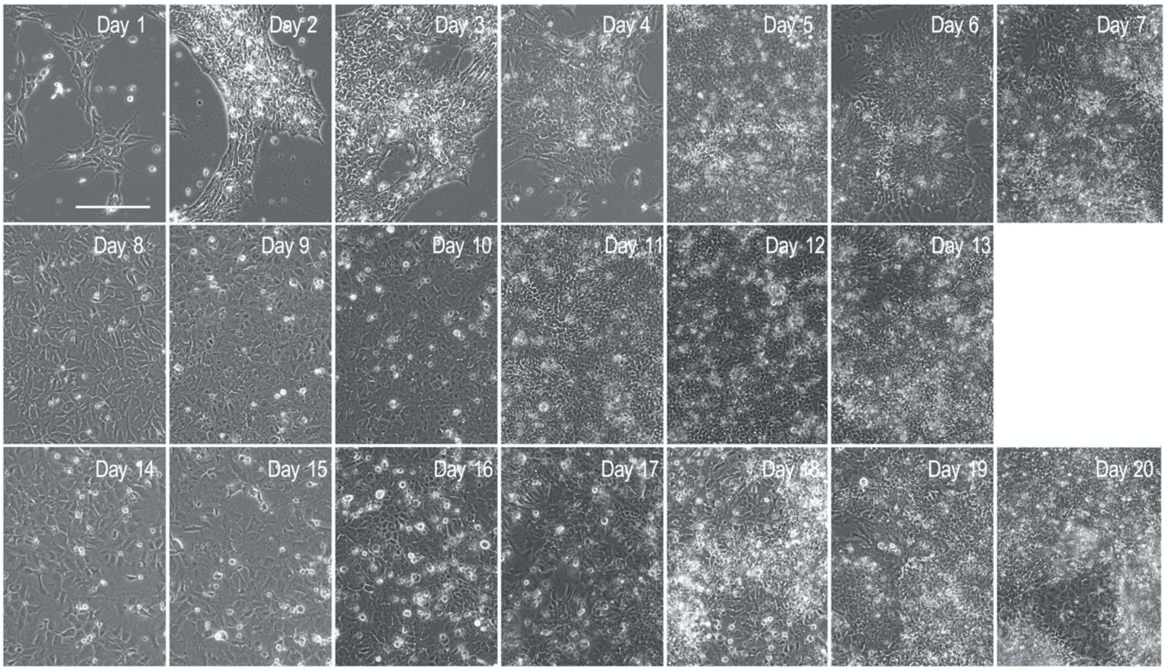


**Supplementary Figure 1.** A timeline of differentiation when the protocol was replicated using H9 hES cells. Representative phase contrast images of cells through differentiation (Days 1 to 20). The images were taken with the EVOS FL microscope, and the scale bar corresponds to 100 µm.

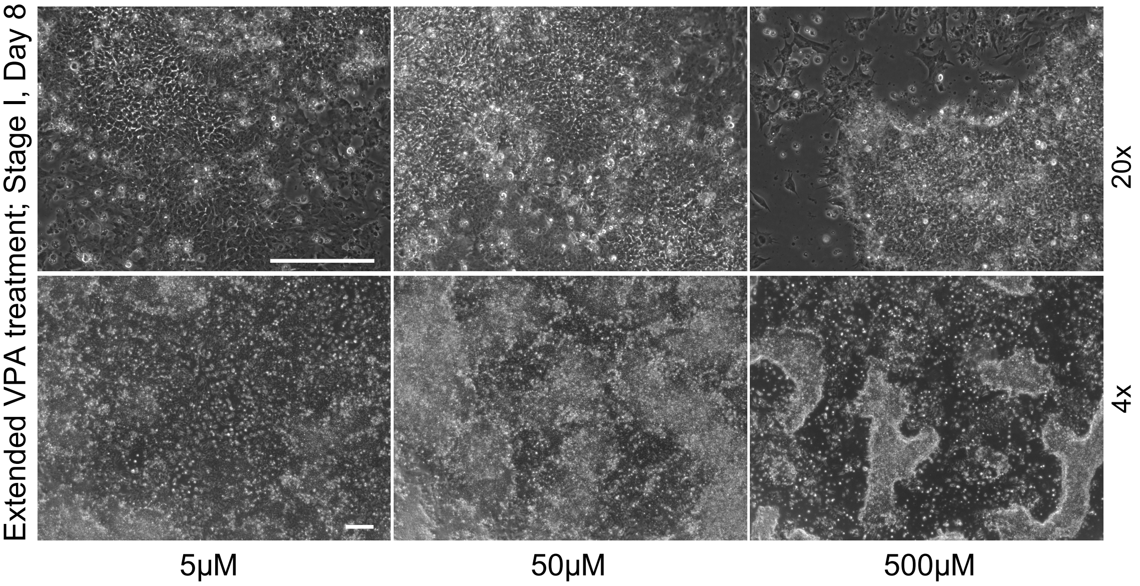


**Supplementary Figure 2.** A panel of the 3 VPA treatment concentrations to demonstrate the morphological differences as seen at day 8 of the extended Stage I cell culture (top row; 20X, bottom row 4X magnification). The lag in differentiation of the cells exposed to 500 µM VPA is more evident after a double washing with 1X PBS to remove debris and dead cells. The images were taken with the EVOS FL microscope, and the scale bar corresponds to 100 µm.
